# *In trans* variant calling reveals enrichment for compound heterozygous variants in genes involved in neuronal development and growth

**DOI:** 10.1101/496133

**Authors:** Allison J. Cox, Fillan Grady, Gabriel Velez, Vinit B. Mahajan, Polly J. Ferguson, Andrew Kitchen, Benjamin W. Darbro, Alexander G. Bassuk

## Abstract

Compound heterozygotes occur when different mutations at the same locus on both maternal and paternal chromosomes produce a recessive trait. Here we present the tool VarCount for the quantification of mutations at the individual level. We used VarCount to characterize compound heterozygous coding variants in patients with epileptic encephalopathy and in the 1000 genomes participants. The Epi4k data contains variants identified by whole exome sequencing in patients with either Lennox-Gastaut Syndrome (LGS) or Infantile Spasms (IS), as well as their parents. We queried the Epi4k dataset (264 trios) and the phased 1000 genomes data (2504 participants) for recessive variants. To assess enrichment, transcript counts were compared between the Epi4k and 1000 genomes participants using minor allele frequency (MAF) cutoffs of 0.5% and 1.0%, and including all ancestries or only probands of European ancestry. In the Epi4k participants, we found enrichment for rare, compound heterozygous mutations in six genes, including three involved in neuronal growth and development – *PRTG* (p=0.00086, 1% MAF, combined ancestries), *TNC* (p=0.0221% MAF, combined ancestries), and *MACF1* (p=0.0245, 0.5% MAF, EU ancestry). Due the total number of transcripts considered in these analyses, the enrichment detected was not significant after correction for multiple testing and higher powered or prospective studies are necessary to validate the candidacy of these genes. However, *PRTG, TNC*, and *MACF1* are potential novel recessive epilepsy genes and our results highlight that compound heterozygous mutations should be considered in sporadic epilepsy.

## Introduction

Using the premise that effective variants are in linkage disequilibrium (LD) with common polymorphisms and haplotypes, linkage and association studies have identified genes involved in the development of traits and pathologies. Upon their identification, the regions flanking associated markers are sequenced to find the linked, penetrant mutation. However, rare variants are often not detectable using LD-based methods. This problem has been alleviated by recent advances in next-generation sequencing (NGS), and the detection of highly penetrant rare variants associated with disease has reduced the heritability gap for such diseases as autism, Crohn’s disease, and osteoporosis (Bomba, Walter, & Soranzo, 2017; Kosmicki, Churchhouse, Rivas, & Neale, 2016). Despite these advances, for most traits and complex disorders the underlying genes and mutations remain elusive.

Recessive traits are caused by mutations in both copies of a gene. The mutations may be homozygous, i.e. identical, or compound heterozygous. Compound heterozygous (CH) mutations are two different mutations in a gene on opposite alleles of a chromosome and it is speculated that compound heterozygous mutations account for many recessive diseases (Li et al., 2010; Sanjak, Long, & Thornton, 2017). Lack of detection of CH may explain a significant portion of missing heritability for all phenotypes (Li et al., 2010; Sanjak et al., 2017; Zhong, Karssen, Kayser, & Liu, 2016). Association studies using polymorphisms are LD-based and recent association studies using rare variants compare total variant burden between cases and controls to account for the contributions of multiple alleles at a locus to phenotype. Importantly, because LD-based studies require recessive mutations to be on the same genetic background and total variant burden analyses are not allele-specific, neither discerns between dominant and recessive models of inheritance.

Burden tests may account for compound heterozygosity if the variants are allocated to one of the two alleles for a gene, i.e. phased. Relatively common variants may be phased assuming linkage to surrounding haplotypes; in families, rare variants are phased using parental genotypes. Once mutations are phased, it may be determined if an individual’s mutations are on different chromosomes, and burden tests that aggregate using an indicator function (i.e. presence of qualifying variants) may assess enrichment for recessive variants.

Here we provide a publicly available tool, VarCount, that is user-friendly and effective for researchers seeking to quantify the presence or absence of a mutation or mutations in a gene at the individual level. VarCount is useful for the quantification of heterozygous, homozygous, or CH mutations per sample. We used VarCount to query the Epi4k (Epi et al., 2013) dataset for rare homozygous and CH mutations and found enrichment for rare, compound heterozygous mutations in six genes, including three involved in neuronal development or growth (*PRTG, TNC*, and *MACF1*). The variants in the 1000 genomes database are now phased (Genomes Project et al., 2010; Genomes Project et al., 2012; Genomes Project et al., 2015), and so genes may be queried for *in trans* combinations of variants. The Epi4k enrichment was identified in comparison to the 1000 genomes participants combining all ancestries and considering only individuals of European ancestry.

## Materials and Methods

### Processing of Epi4k vcf files

The Epi4k data (Epi et al., 2013) were accessed by permission via the Database of Genotypes and Phenotypes (dbGaP Study Accession, phs000653.v2.p1). Individual vcf files were combined using the CombineVariants function in GATK (McKenna et al., 2010). The vcf files were then annotated with minor allele frequencies (MAFs) from EVS (Exome Variant Server, NHLBI GO Exome Sequencing Project (ESP), Seattle, WA (URL: http://evs.gs.washington.edu/EVS/)), 1000 genomes and ExAC (Monkol Lek et al., 2015), and with information regarding the effect of each variant using SNPSift/SNPEff (Cingolani et al., 2012). The databases used for annotation were dbNSFP2.9 (for MAF and CADD score) and GRCh37.75 for protein effect prediction. SnpSift was used to remove any variants not inducing a protein-changing event (not “HIGH” or “MODERATE” impact) based on SNPEff annotation – this includes missense, nonsense, splice-site and insertion/deletion variants. Variants with quality flags and multiallelic variants, i.e. those with more than two known nucleotide values, were also removed. Variants remaining after filtering were cross-referenced with the 1000 genomes variants from the same MAF threshold to ensure that any variants removed from one dataset were removed from the other. The annotated vcf was used as input for VarCount. Ancestry for each exome was determined using LASER (Wang et al.) and this information was input to VarCount via the SampleInfo.txt file. Ancestry and phenotype information for each proband are described in Supplementary Table 1. In addition to the annotated vcf file, the parameters.txt and subjectinfo.txt (containing sex and ancestry information) were used as input. Within the parameters file, the following qualifications were selected: (1) counting at the transcript (rather than gene) level, (2) protein-changing effects, (3) MAF threshold of either 0.005 or 0.01, (4) all within-dataset and annotated (1000 genomes, ExAC and EVS) MAFs, and (5) either compound heterozygous or homozygous mutations. Analyses were run separately for the two MAFs and using all Epi4k probands (264) and only those of Eurpoean ancestry (207). Because the variants were not phased, VarCount was used to query the vcf file for individuals with two or more mutations in each transcript. The output, a list of counts for each transcript was then used to query the parental vcf files for genotype information to determine which sets of variants composed *in trans* combinations of mutations. Final counts were determined using parental genotype information. Custom python scripts were used to query for parental genotypes and to count true compound heterozygotes or homozygotes. *De novo* mutations were excluded in the determination of true *in trans* mutations.

### Processing of 1000 genomes vcf files

Vcf files for the 2504 participants in the 1000 genomes sequencing project (Genomes Project et al., 2015) were downloaded by chromosome from the 1000 genomes ftp site. To reduce input file size, the genomic regions for the hg19 mRNA transcripts were downloaded via UCSC’s Table Browser and used to remove non-coding regions from the vcf files. Including all exons from UCSC allowed for a more conservative analysis, given that the Epi4k data were sequenced using various exome captures, which are not inclusive of all possible exons. The variants were annotated and filtered via the same steps as the Epi4k vcf file. Multi-allelic variants were also removed prior to analysis by VarCount. A diagram showing the steps involved in processing and analyzing the variant files is shown in (Figure 1).

**Figure 1.**
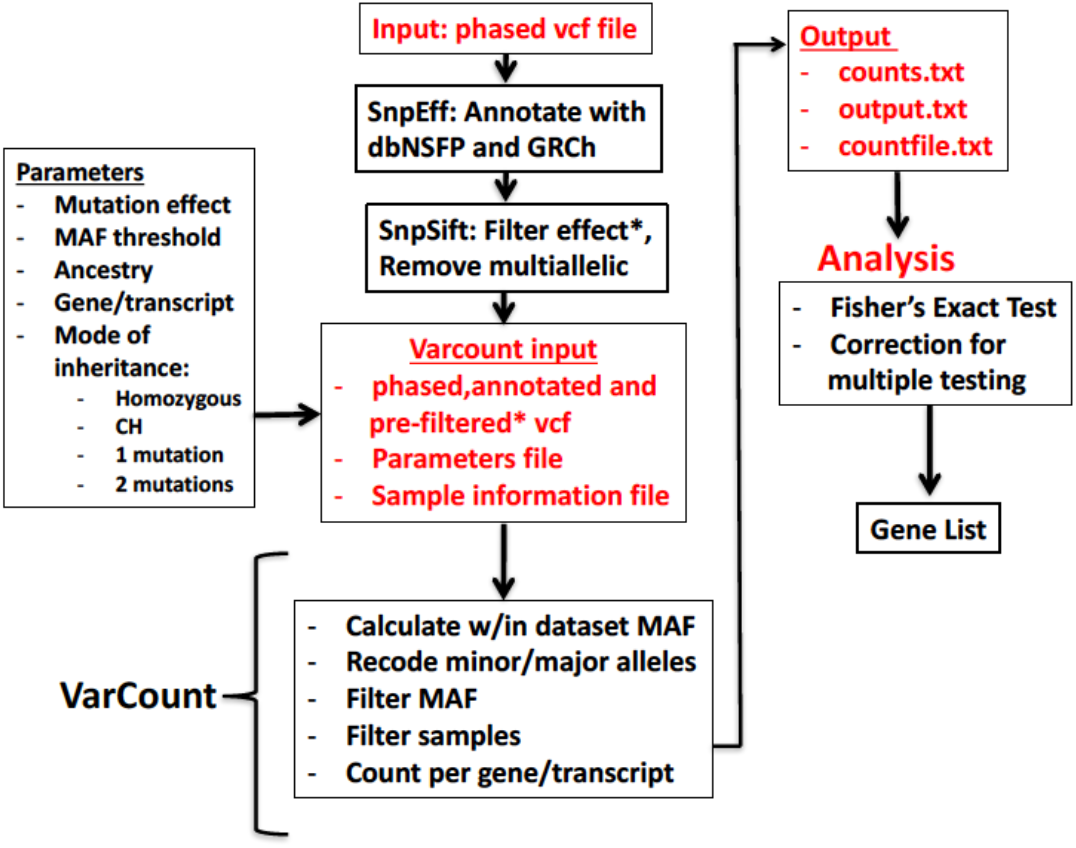
Flow diagram for the processing and analysis of variant lists. Vcf files are annotated and filtered using SnpSift/SnpEff. Final vcf along with parameter and sample information files are input to VarCount. The imput files are processed to recode minor and major alleles when the MAF > 0.5 and to count the number of individuals with mutations qualifying based on information in the parameter file. The final output lists for every transcript or gene, the number of individuals with qualified mutations in that locus (counts.text), which individuals have the mutation(s) (countfile.txt), and which mutations are harbored by each individual (output.txt).

VCF files were queried for homozygous and compound heterozygous mutations using VarCount. Because the variants in the 1000 genomes vcf files are phased, determining true compound heterozygotes is automatic using VarCount. In addition to the annotated vcf file, the parameters.txt and subjectinfo.txt (containing sex and ancestry information) were used as input. Within the parameters file, the same qualifications used in the Epi4k analysis were selected: Analyses were run for each of the two MAFs (0.5% and 1.0%) and for all 1000 genomes participants and using only those of European (EUR) ancestry. The final output from analyses was for each MAF cutoff and for each population, counts for every transcript in which at least one individual harbored recessive variants.

### Epi4k statistical analysis

Using R statistical software, a Fisher’s exact test was used to detect transcripts with significant differences in the proportion of individuals with homozygous or compound heterozygous variants between the Epi4k dataset and the 1000 genomes dataset. Odds ratios and p-values were calculated using the number of individuals with and without qualifying mutations in each superpopulation. Analyses were performed using all ancestries, and for only individuals of European ancestry. Both Bonferroni and Benjamini-Hochberg adjustments were used to determine significance thresholds after correction for multiple testing. The number of tests was based on the number of transcripts with at least one individual in either the Epi4k or 1000 genomes dataset with *in trans* coding variants with minor allele frequencies below the set threshold.

### Structural modeling of PRTG

The three-dimensional structure of Protogenin (PRTG) was modeled off the crystal structure of the human receptor protein tyrosine phosphatase sigma (PDB: 4PBX; 25.1% sequence identity) using MODELLER 9.14 (Webb & Sali, 2016). The resultant model superimposed with the template had an RSMD of 4.94 Å over 442 Ca atoms. Charges and hydrogen atoms were added to the wild-type and mutant FGR models using PDB2PQR (Dolinsky, Nielsen, McCammon, & Baker, 2004). Electrostatic potentials were calculated using APBS (Konecny, Baker, & McCammon, 2012) as described previously (Cox et al., 2017; Moshfegh et al., 2016; Toral et al., 2017). Protein and solvent dielectric constants were set to 2.0 and 78.0, respectively. All structural figures were generated by PyMOL(W).

## Results

### Varcount: Mutation Quantification at the Individual level

Varcount is a free, open source tool useful for the quantification of heterozygous, homozygous, or compound heterozygous mutations per sample. Input variants may be phased or unphased. All python scripts and supporting files may be downloaded from Github at https://github.com/GeneSleuth/VarCount. Supporting files include the “parameters.txt” file where the user may select variant filters for mutation effect, minor allele frequency, and inheritance pattern (homozygous, compound heterozygous, one mutation or two mutations), and Sample filters based on information entered into the “SampleInfo file”. Input vcf files must be annotated with SNPSift/SNPEff using the dbNSFP and GRCh37/38 databases. A readme file with instructions is also provided. A flow diagram with the steps involved in processing of data is depicted in Figure 1.

### Compound heterozygous mutations in Epi4k probands reveal novel epilepsy genes

We used VarCount to query the Epi4k dataset for rare homozygous and CH mutations. The Epi4k data are whole exome data from 264 trios with a child affected by epileptic encephalopathy, either infantile spasms (IS) or Lennox-Gastaut syndrome (LGS) (6). Counts were performed using individuals of all ancestries or just those of European ancestry (207/264). Individuals from the 1000 genomes study were used as controls. The individual counts and p-values for the analyses are listed in (Supplementary Tables 2-5). Including only rare variants (MAFs below 0.5% and 1.0%) determined enrichment for compound heterozygous mutations in six genes. For combined ancestries, the six genes are in order of significance: *OSBP2, PRTG, ABCC11, MACF1, STAB1*, and *TNC. PRTG* and *TNC* were also highly ranked in the 1% MAF analysis, with one additional count for each transcript. Variants for all six genes are listed in Table 1. In our analysis of just individuals of European ancestry, *MACF1* was the most significantly enriched gene using a 0.5% MAF. The p-values indicated in Table 1 are for individual tests; there were no p-values significant after correction for multiple testing.

**Table 1.**
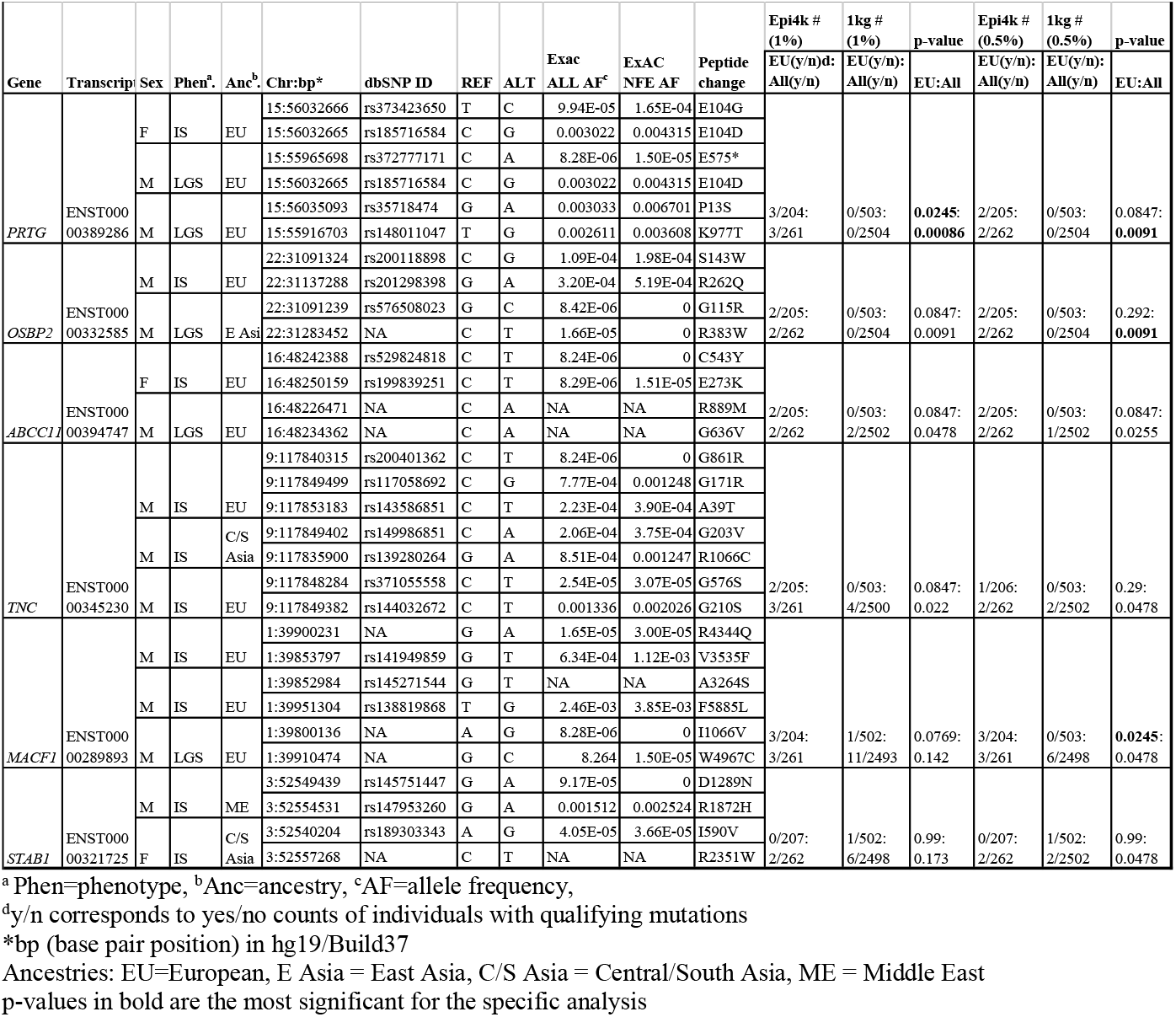
Rare (< 0.5% and 1.0% MAF) Compound Heterozygous Mutations in Epi4k participants.

The variants for the three individuals with CH *PRTG* mutations are depicted in Figure 2A. Because of the concentration of mutations at position E104, we performed structural modeling to predict the pathogenicity of the PRTG mutations. The p.Glu104Gly and p.Glu104Asp mutations localize to the immunoglobulin (Ig)-like domain 1 (Fig. 2A). Ig-like domains are responsible for mediating protein-protein and protein-peptide interactions. The p.Glu104Gly disrupts a negative charge in the Ig-like 1 domain. This loss of charge may disrupt interactions with putative PRTG-binding partners (Fig. 2B).

**Figure 2.**
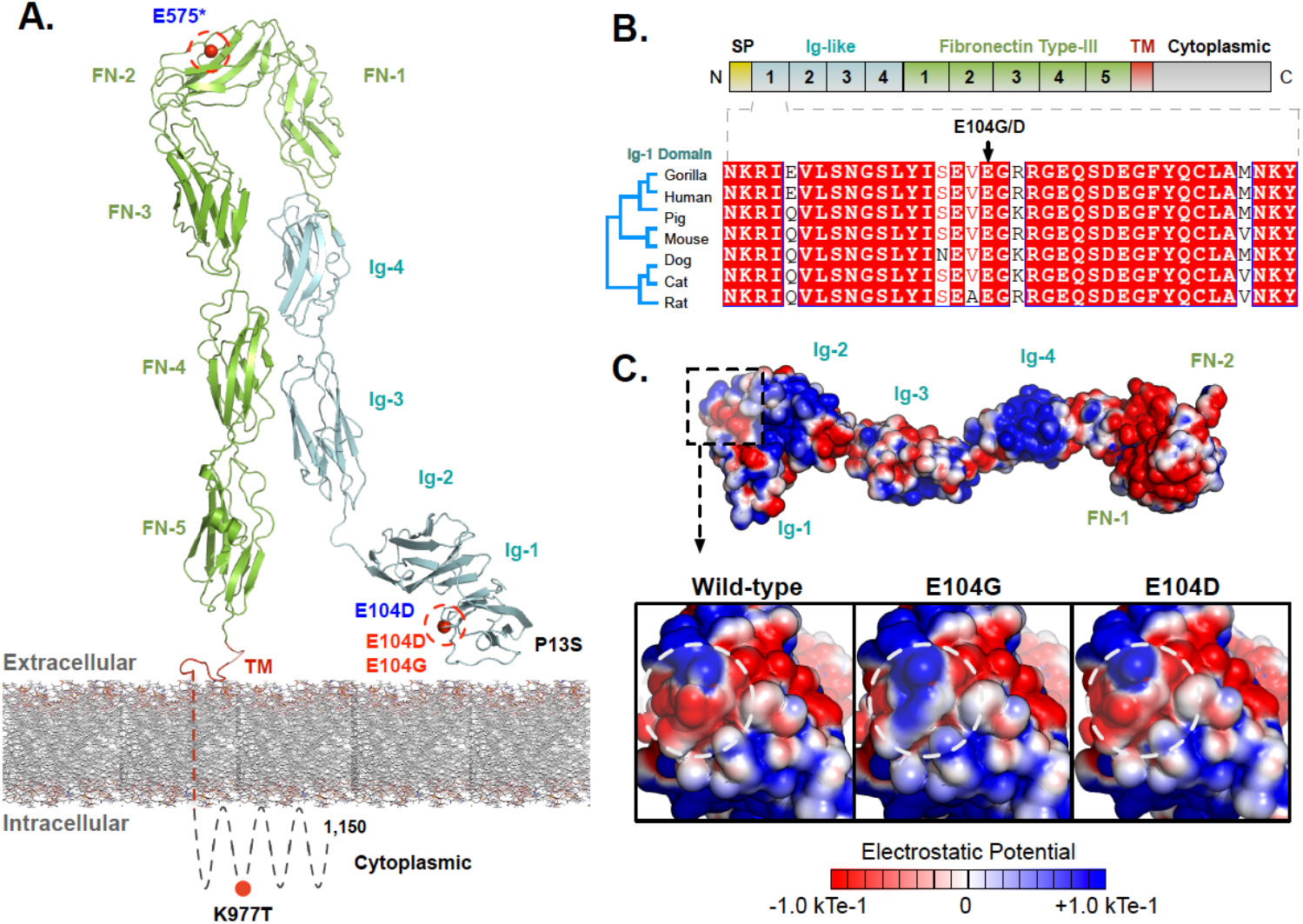
PRTG compound heterozygous mutations in Epi4k probands. (A) Protein diagram indicating mutation locations in each child. The three pairs of *in trans* mutations are indicated in red, blue and black. The mutations in red and blue were found using a 0.5% MAF threshold and the mutations in black were found using a 1% MAF threshold. (B) Bottom: electrostatic potential surface of PRTG calculated in APBS. Top: Close-up of the PRTG electrostatic potential surface at the site of mutation. The p.Glu104Gly mutation leads to a loss of negative charge, which may disrupt interactions with putative PRTG binding partners. The p.Glu104Asp mutation does not lead to a change in charge or electrostatic potential.

The *de novo* variants identified by Epi4K Consortium and the Epilepsy Phenome/Genome Project (Epi et al., 2013) in the nine probands with either *PRTG, TNC*, or *MACF1* recessive variants are described in Table 2. For the three patients with compound heterozygous *PRTG* variants, one patient harbors a *de novo* missense variant in *HSF2*, the second has a nonsense variant in *CELSR1*, and the third patient has two *de novo* variants – a missense in Fam102A and a 3’UTR variant in *USP42. De novo* mutations were only reported in one of the probands with *in trans TNC* variants – a missense variant in *DIP2C* and a splice donor change in *IFT172*. All three patients with compound heterozygous variants in *MACF1* were reported to have *de novo* mutations. The first patient has a 5’ and 3’UTR *de novo* variant in *FAM19A2* and *GLRA2*, respectively and the second patient also has a 3’UTR *de novo* change in the gene *LRRC8D* and a missense change in *SNX30*. One *de novo* variant was identified in the third proband in the gene *FAM227A*. Polyphen2 categories and CADD scores for each *de novo* variant as well as missense and loss-of-function constraint metric values for each gene (from ExAC) are also listed in Table 2. The z-score is a ratio of expected to identified missense variants in a particular gene, and pLI is a gene’s probability of being loss-of-function intolerant. These constraint metrics are calculated using genomic data from controls without severe genetic diseases in the ExAC database (Monkol Lek et al., 2015).

**Table 2.**
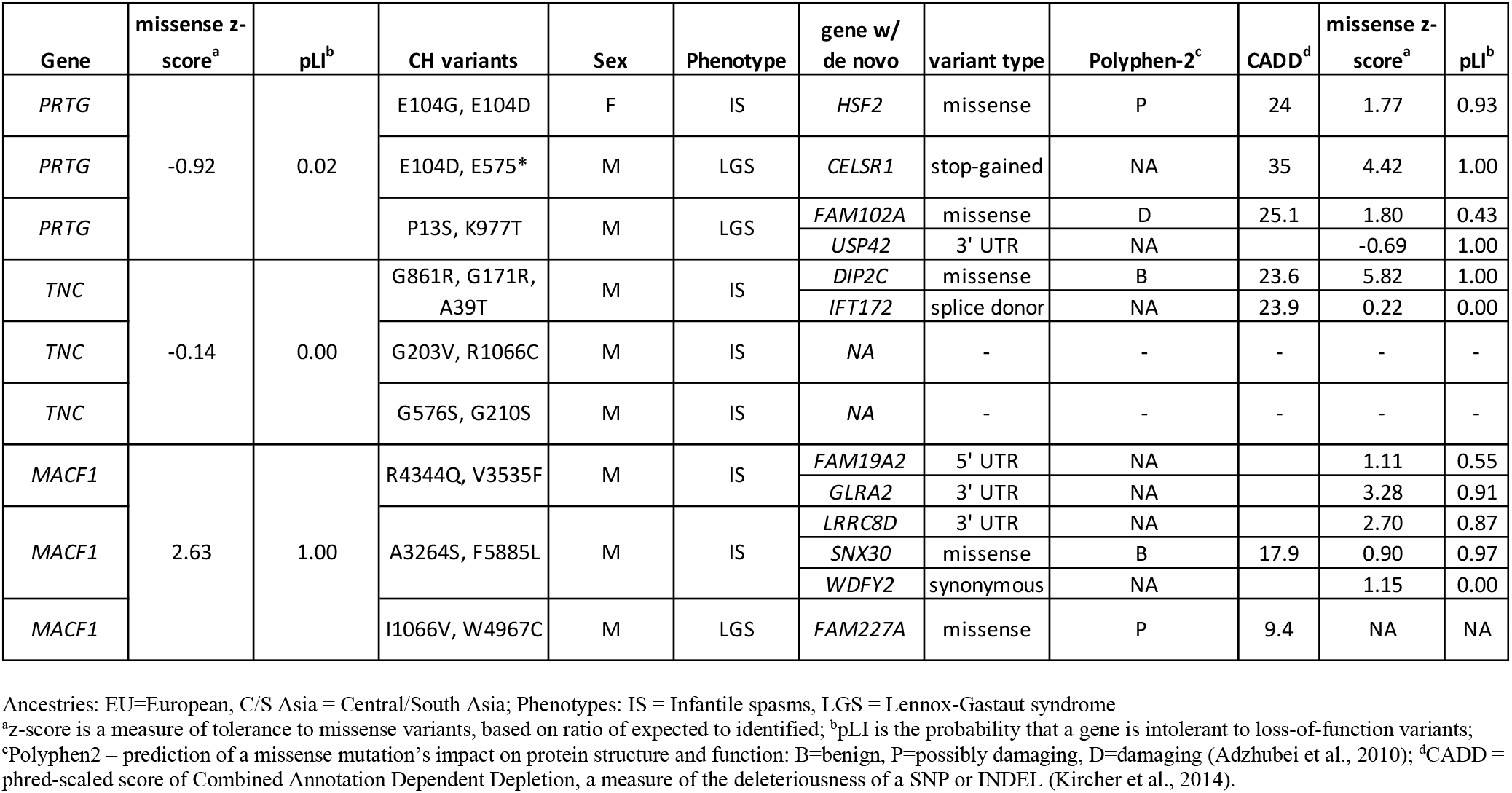
*De novo* variants in Epi4k probands with CH variants in PRTG, TNC, or MACF1.

## Discussion

Epileptic encephalopathies are a group of severe, early-onset seizure disorders with consistent EEG abnormalities that over time interfere with development and cause cognitive decline (Covanis, 2012). The Epi4k dataset contains exome sequence from 264 trios that include a proband with epileptic encephalopathy, either Lennox-Gastaut Syndrome (LGS) or Infantile Spasms (IS). LGS is characterized by frequent, mixed epileptic seizures that arise most frequently between the ages of 3 and 5 (Amrutkar and Riel-Romero, 2018). IS occurs during the first year of life and is cryptic in its presentation, with mild head bobbing and is often not detected until the seizures have caused significant neurological damage (Kossoff, 2010). IS often progress into LGS over time.

We developed a free and user-friendly tool, VarCount, to query vcf files for individuals harboring variants that qualify according to user specification. To test its function, we used VarCount to quantify rare, compound heterozygous mutations in probands from the Epi4k trio dataset and found enrichment for variants in six genes including *PRTG, TNC* and *MACF1. PRTG* codes for protogenin, a member of the immunoglobulin superfamily that is involved in axis elongation and neuronal growth during early vertebrate development (Toyoda, Nakamura, & Watanabe, 2005; Vesque, Anselme, Couve, Charnay, & Schneider-Maunoury, 2006). TNC and MACF1 are also directly involved in neuronal development and/or growth. TNC (Tenascin-C) is an extracellular matrix glycoprotein involved in axonal growth and guidance (Jakovcevski, Miljkovic, Schachner, & Andjus, 2013). Seizures up-regulate *TNC* in the hippocampus, and in a pilocarpine epilepsy model up-regulation was shown to be mediated by TGF-β signalling (Mercado-Gomez, Landgrave-Gomez, Arriaga-Avila, Nebreda-Corona, & Guevara-Guzman, 2014). MACF1 is a cytoskeletal crosslinking protein highly expressed in the brain and is crucial for neuron development and migration (Moffat, Ka, Jung, Smith, & Kim, 2017). *MACF1* mutations are associated with the neurological pathologies Parkinson’s disease, autism, and schizophrenia (Moffat et al., 2017). Recently, highly penetrant *de novo MACF1* mutations were identified in several patients with a newly characterized lissencephaly with a complex brain malformation (Dobyns et al., 2018). This new phenotype highlights *MACF1* mutations’ variable impact on disease pathogenesis. Given both the enrichment in Epi4k probands for compound heterozygous mutations in these genes as well as their known involvement in neuronal processes, we suggest that *PRTG, TNC*, and *MACF1* are candidate recessive epilepsy genes.

The primary publication reporting analysis of the Epi4k trio dataset was a description of *de novo* mutations in the probands (Epi et al., 2013). An analysis of compound heterozygous variants was also reported, using a minor allele frequency cutoff of 0.15%, which is lower than the cutoff used in the work presented here. In this analysis, the parents were used as internal controls, and compound heterozygous variants in 351 genes were identified, without genome-wide significance. The authors only listed five of the genes which are known to cause Mendelian disorders that include a seizure phenotype – *ASPM, CNTNAP2, GPR98, PCNT*, and *POMGNT1*. In our analysis using the 1000 genomes participants as controls, enrichment for compound heterozygous variants was not detected in any of these genes. Using the number of individuals with *in trans* variants in a gene (transcript) as an indicator function required at least two probands to have qualifying variants in order to detect single-test significance, with complete absence of qualifying variants in controls. It is clear from the analyses using either internal controls or the 1000 genomes as controls that a larger sample size is required to achieve genome-wide significance.

The *de novo* variants reported by the Epi4K Consortium and the Epilepsy Phenome/Genome Project (Epi et al., 2013) in the nine probands with compound heterozygous variants in *PRTG, TNC*, or *MACF1* are described in Table 2. Of the twelve genes with *de novo* variants identified in the nine patients, three are implicated in neurological disease. *CELSR1* is a planar cell polarity gene in which mutations are known to cause neural tube defects including spina bifida (Robinson et al., 2012). *De novo* deletions of *DIP2C* have been reported in two patients with cerebral palsy, one of whom also had ADHD, and the other had seizures in infancy (Zarrei et al., 2018). In another report, deletions including *DIP2C* and/or *ZMYND11* were identified in several patients with developmental delay including three patients with seizures (DeScipio et al., 2012). GLRA2 is a glycine receptor involved in neurodevelopment in which mutations are implicated in autism, (Pilorge et al., 2016; Lin et al., 2017) including a patient with comorbid epilepsy (Zhang et al., 2017).

Of the *de novo* variants reported in these genes, the nonsense variant in *CELSR1* identified in one of the probands with in trans *PRTG* variants is the most likely to be pathogenic. However, regarding their involvement in neural tube defects, mutations in *CELSR1* are thought to contribute to pathogenesis but not in a Mendelian fashion, as variants have been found to be inherited from unaffected parents or to be ineffective in functional assays (Robinson et al., 2012; Allache et al., 2012). The nonsense *CELSR1* mutation in the patient reported here may contribute to epilepsy in the presence of a genetic modifier. The *de novo* missense mutation in *DIP2C* is predicted to be deleterious (CADD = 23.6) and has a low rate of benign missense variation based on constraint metrics (z = 5.82). The *de novo* variant in *GLRA2* is in the 3’UTR so it is difficult to predict its impact on gene function and subsequent pathogenicity.

The compound heterozygous variants in *PRTG, TNC*, and *MACF1* are similarly variable in predicted pathogenicity, with CADD scores ranging from between less than one to thirty-eight. *PRTG* and *TNC* both have constraint metrics indicative of a high tolerance to both missense and loss-of-function variants, while *MACF1* is moderately intolerant of missense variants (z = 2.63) and extremely intolerant of loss-of-function variants (pLI = 1.0). Interestingly, aside from the 3’UTR variant in *GLRA2*, none of the *de novo* variants in the Epi4k participants with *MACF1* compound heterozygous variants are in genes associated with neurological disease or predicted with confidence to have a negative impact on gene function. This, in addition to *MACF1*’s intolerance to missense or nonsense variants, is supportive of the pathogenicity of the biallelic variants in the gene.

In summary, we present a free tool VarCount for the quantification of qualifying mutations as an indicator function per individual in the analysis of variant lists (vcf files). We used VarCount to assess enrichment of rare, coding, compound heterozygous variants in a cohort of 264 epilepsy probands and found enrichment in three genes involved in neurodevelopmental processes – *PRTG, TNC* and *MACF1*. A missense change at the E104 residue of PRTG was identified three times in two different probands. Significance was not maintained after correction for multiple testing, and larger cohorts or candidate gene studies using a different sample set are necessary to validate this enrichment. In the context of the *de novo* mutations also present in these patients, experimentation is necessary in order to delineate if the compound heterozygous or *de novo* mutations, or both, are pathogenic in the development of epileptic encephalopathy. *PRTG, TNC*, and *MACF1* are candidate recessive epilepsy genes and our work highlights that inheritance of compound heterozygous variants should not be excluded from gene discovery or diagnostic analyses of patients with epilepsy.

## ACKNOWLEDGEMENTS

This work was supported by the following grants: T32GM008629 (XX), T32GM082729-01(XX), T32GM007337 (XX and XX), R01AR059703 (PJF, VBM and AGB) and R01NS098590 (AGB, PJF).

---

**Supplementary Table 1:**
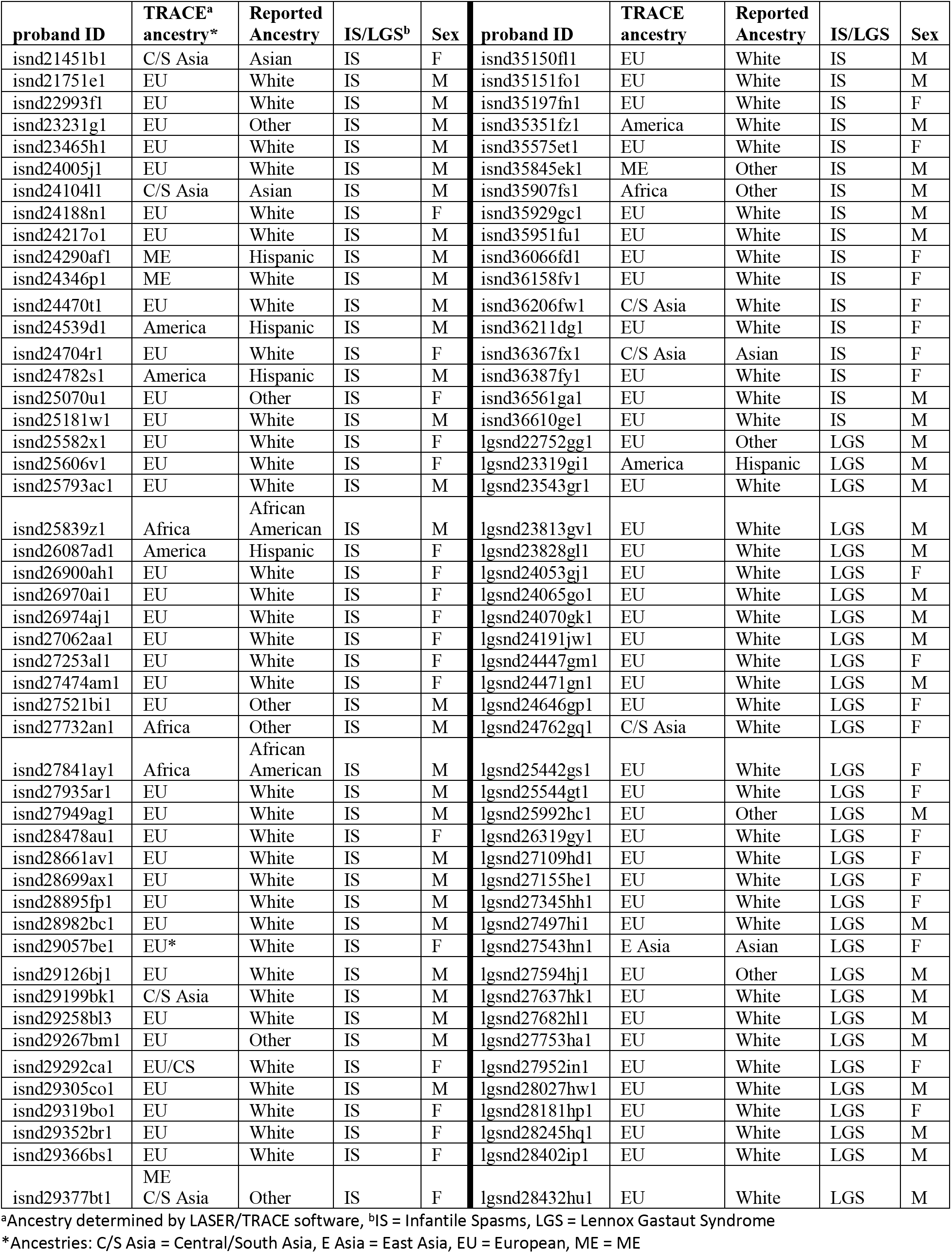
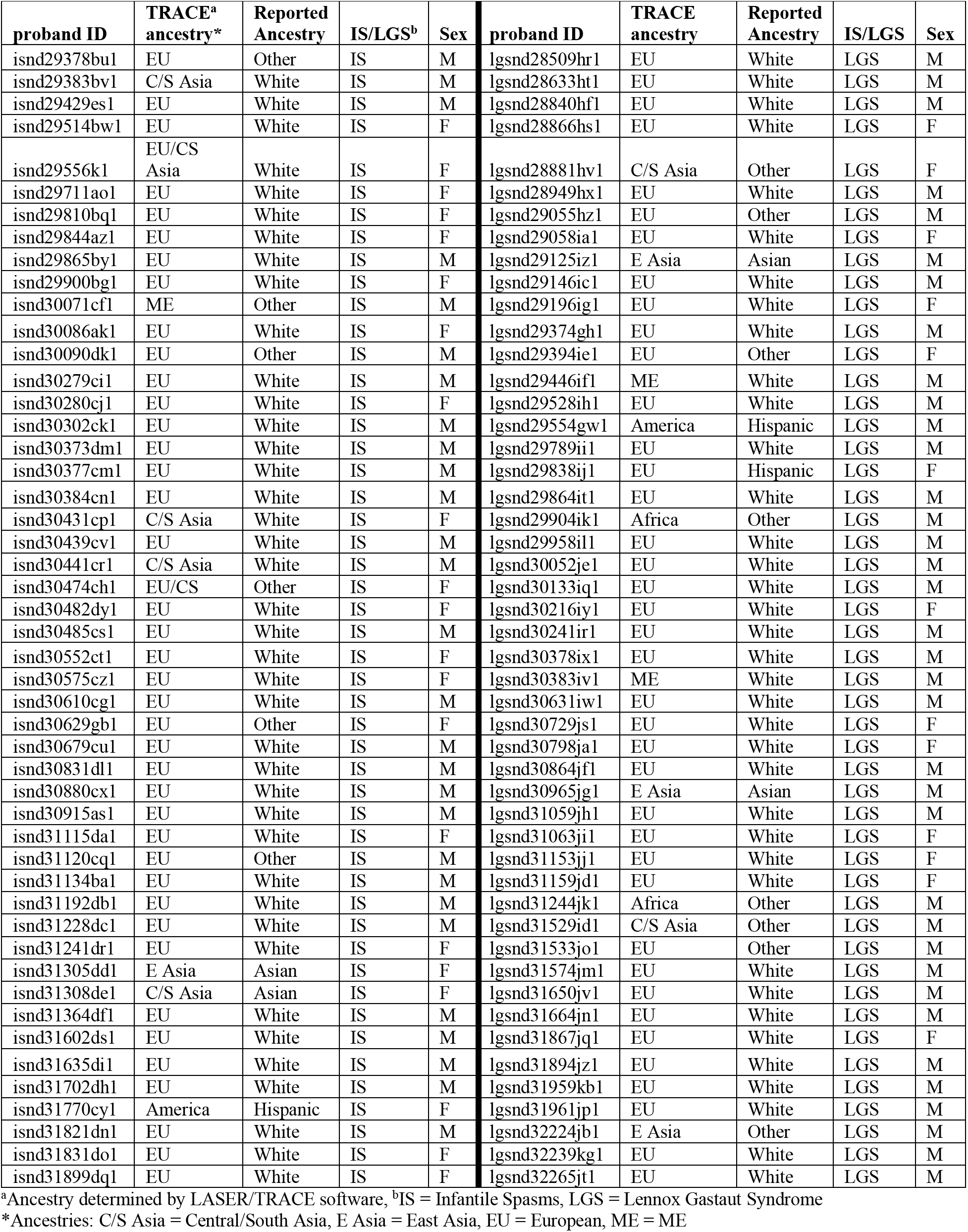
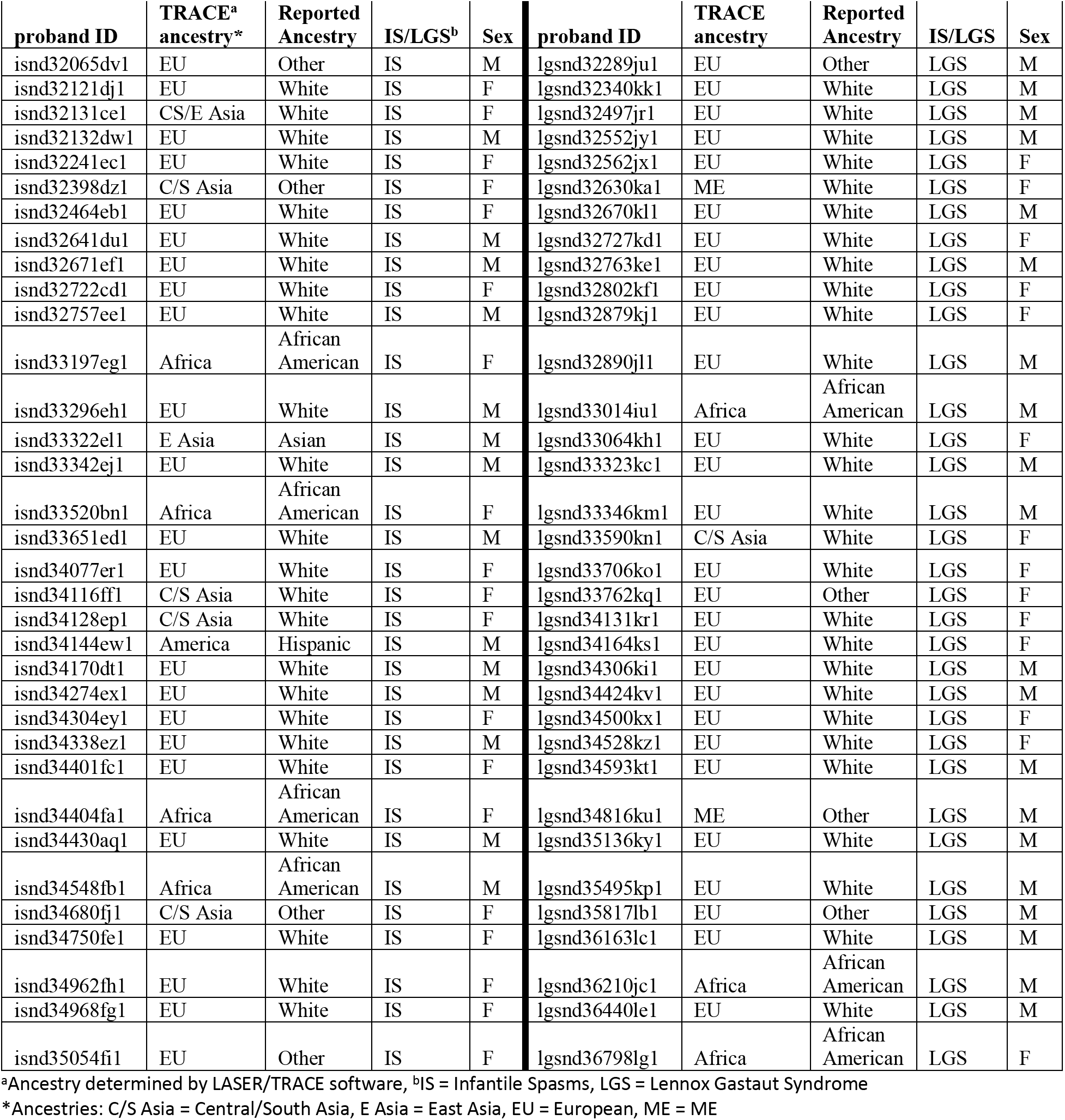
Ancestry and Phenotype Information for Epi4k probands.

**Supplementary Table 2.**
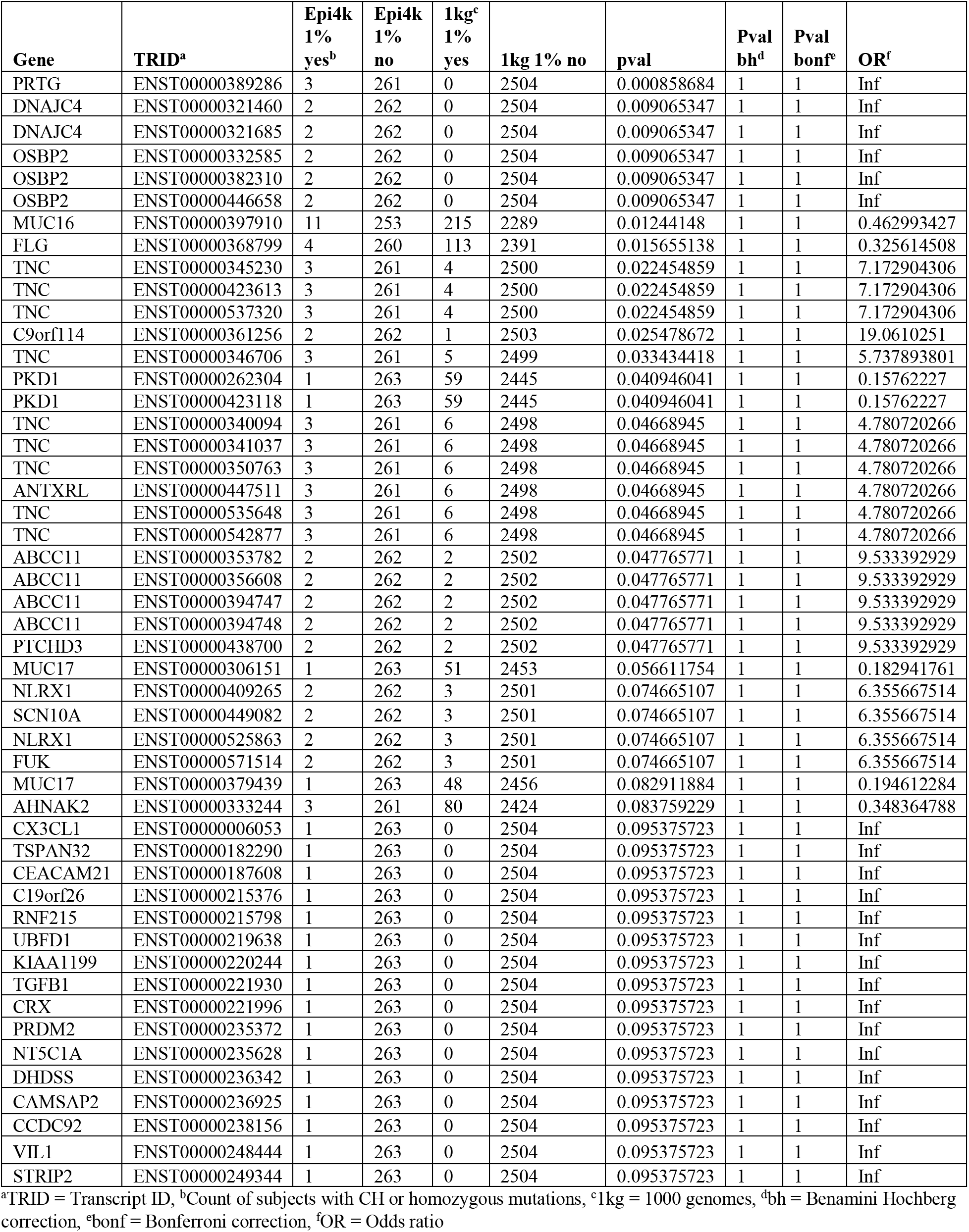
Epi4k and 1000 genomes CH and homozygous counts and p-values, all ancestries, 1%MAF, top 50.

**Supplementary Table 3.**
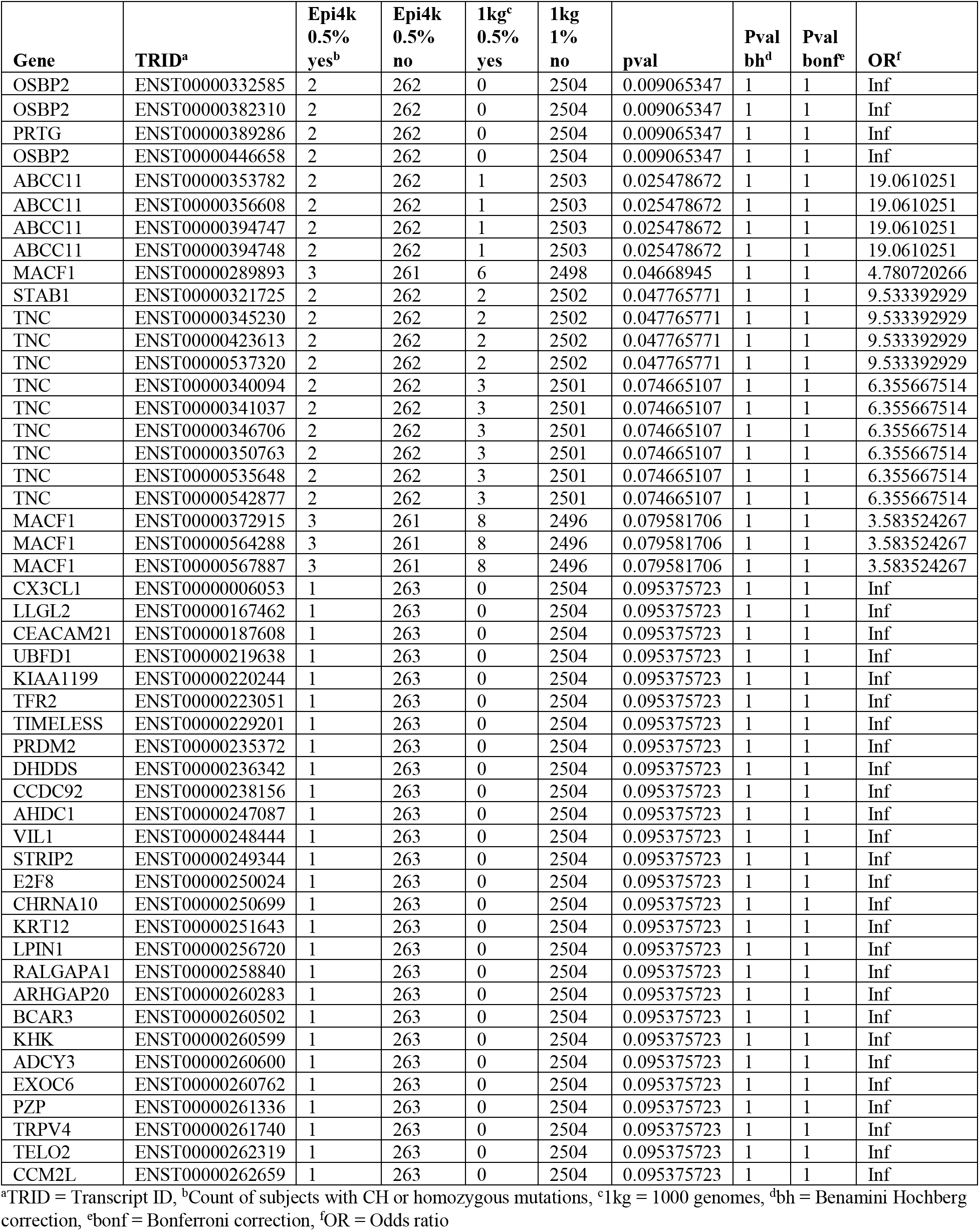
Epi4k and 1000 genomes CH and homozygous counts and p-values, all ancestries, 0.5%MAF, top 50.

**Supplementary Table 4.**
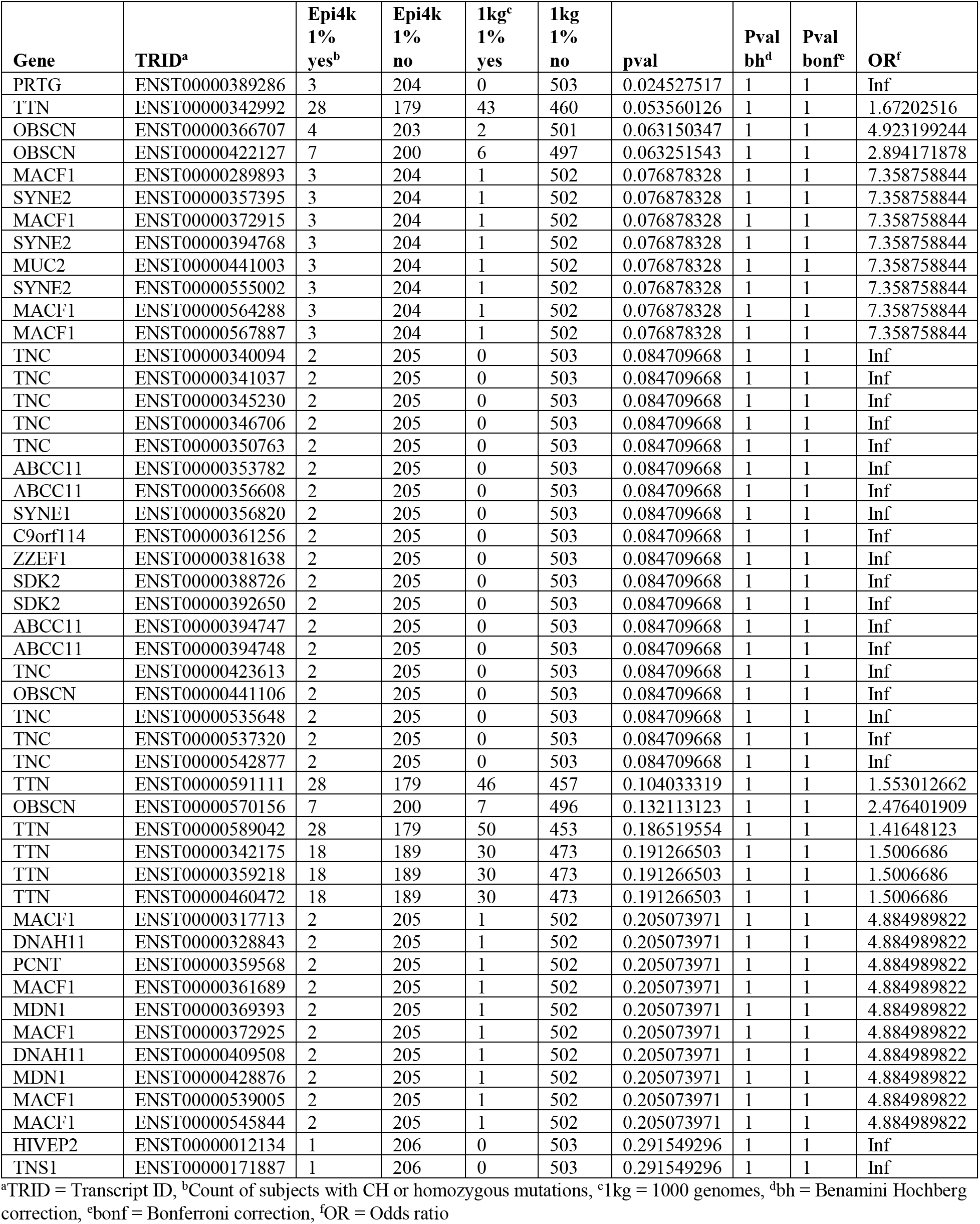
Epi4k and 1000 genomes CH and homozygous counts and p-values, EU ancestry, 1%MAF, top 50.

| Gene | TRID<sup>a</sup> | Epi4k<br>1%<br>yes<sup>b</sup> | Epi4k<br>1%<br>no | 1kg<sup>c</sup><br>1%<br>yes | 1kg<br>1%<br>no | pval | Pval<br>bh<sup>d</sup> | Pval<br>bonf<sup>e</sup> | OR<sup>f</sup> |
| --- | --- | --- | --- | --- | --- | --- | --- | --- | --- |
| PRTG | ENST00000389286 | 3 | 204 | 0 | 503 | 0.024527517 | 1 | 1 | Inf |
| TTN | ENST00000342992 | 28 | 179 | 43 | 460 | 0.053560126 | 1 | 1 | 1.67202516 |
| OBSCN | ENST00000366707 | 4 | 203 | 2 | 501 | 0.063150347 | 1 | 1 | 4.923199244 |
| OBSCN | ENST00000422127 | 7 | 200 | 6 | 497 | 0.063251543 | 1 | 1 | 2.894171878 |
| MACF1 | ENST00000289893 | 3 | 204 | 1 | 502 | 0.076878328 | 1 | 1 | 7.358758844 |
| SYNE2 | ENST00000357395 | 3 | 204 | 1 | 502 | 0.076878328 | 1 | 1 | 7.358758844 |
| MACF1 | ENST00000372915 | 3 | 204 | 1 | 502 | 0.076878328 | 1 | 1 | 7.358758844 |
| SYNE2 | ENST00000394768 | 3 | 204 | 1 | 502 | 0.076878328 | 1 | 1 | 7.358758844 |
| MUC2 | ENST00000441003 | 3 | 204 | 1 | 502 | 0.076878328 | 1 | 1 | 7.358758844 |
| SYNE2 | ENST00000555002 | 3 | 204 | 1 | 502 | 0.076878328 | 1 | 1 | 7.358758844 |
| MACF1 | ENST00000564288 | 3 | 204 | 1 | 502 | 0.076878328 | 1 | 1 | 7.358758844 |
| MACF1 | ENST00000567887 | 3 | 204 | 1 | 502 | 0.076878328 | 1 | 1 | 7.358758844 |
| TNC | ENST00000340094 | 2 | 205 | 0 | 503 | 0.084709668 | 1 | 1 | Inf |
| TNC | ENST00000341037 | 2 | 205 | 0 | 503 | 0.084709668 | 1 | 1 | Inf |
| TNC | ENST00000345230 | 2 | 205 | 0 | 503 | 0.084709668 | 1 | 1 | Inf |
| TNC | ENST00000346706 | 2 | 205 | 0 | 503 | 0.084709668 | 1 | 1 | Inf |
| TNC | ENST00000350763 | 2 | 205 | 0 | 503 | 0.084709668 | 1 | 1 | Inf |
| ABCC11 | ENST00000353782 | 2 | 205 | 0 | 503 | 0.084709668 | 1 | 1 | Inf |
| ABCC11 | ENST00000356608 | 2 | 205 | 0 | 503 | 0.084709668 | 1 | 1 | Inf |
| SYNE1 | ENST00000356820 | 2 | 205 | 0 | 503 | 0.084709668 | 1 | 1 | Inf |
| C9orf114 | ENST00000361256 | 2 | 205 | 0 | 503 | 0.084709668 | 1 | 1 | Inf |
| ZZEF1 | ENST00000381638 | 2 | 205 | 0 | 503 | 0.084709668 | 1 | 1 | Inf |
| SDK2 | ENST00000388726 | 2 | 205 | 0 | 503 | 0.084709668 | 1 | 1 | Inf |
| SDK2 | ENST00000392650 | 2 | 205 | 0 | 503 | 0.084709668 | 1 | 1 | Inf |
| ABCC11 | ENST00000394747 | 2 | 205 | 0 | 503 | 0.084709668 | 1 | 1 | Inf |
| ABCC11 | ENST00000394748 | 2 | 205 | 0 | 503 | 0.084709668 | 1 | 1 | Inf |
| TNC | ENST00000423613 | 2 | 205 | 0 | 503 | 0.084709668 | 1 | 1 | Inf |
| OBSCN | ENST00000441106 | 2 | 205 | 0 | 503 | 0.084709668 | 1 | 1 | Inf |
| TNC | ENST00000535648 | 2 | 205 | 0 | 503 | 0.084709668 | 1 | 1 | Inf |
| TNC | ENST00000537320 | 2 | 205 | 0 | 503 | 0.084709668 | 1 | 1 | Inf |
| TNC | ENST00000542877 | 2 | 205 | 0 | 503 | 0.084709668 | 1 | 1 | Inf |
| TTN | ENST00000591111 | 28 | 179 | 46 | 457 | 0.104033319 | 1 | 1 | 1.553012662 |
| OBSCN | ENST00000570156 | 7 | 200 | 7 | 496 | 0.132113123 | 1 | 1 | 2.476401909 |
| TTN | ENST00000589042 | 28 | 179 | 50 | 453 | 0.186519554 | 1 | 1 | 1.41648123 |
| TTN | ENST00000342175 | 18 | 189 | 30 | 473 | 0.191266503 | 1 | 1 | 1.5006686 |
| TTN | ENST00000359218 | 18 | 189 | 30 | 473 | 0.191266503 | 1 | 1 | 1.5006686 |
| TTN | ENST00000460472 | 18 | 189 | 30 | 473 | 0.191266503 | 1 | 1 | 1.5006686 |
| MACF1 | ENST00000317713 | 2 | 205 | 1 | 502 | 0.205073971 | 1 | 1 | 4.884989822 |
| DNAH11 | ENST00000328843 | 2 | 205 | 1 | 502 | 0.205073971 | 1 | 1 | 4.884989822 |
| PCNT | ENST00000359568 | 2 | 205 | 1 | 502 | 0.205073971 | 1 | 1 | 4.884989822 |
| MACF1 | ENST00000361689 | 2 | 205 | 1 | 502 | 0.205073971 | 1 | 1 | 4.884989822 |
| MDN1 | ENST00000369393 | 2 | 205 | 1 | 502 | 0.205073971 | 1 | 1 | 4.884989822 |
| MACF1 | ENST00000372925 | 2 | 205 | 1 | 502 | 0.205073971 | 1 | 1 | 4.884989822 |
| DNAH11 | ENST00000409508 | 2 | 205 | 1 | 502 | 0.205073971 | 1 | 1 | 4.884989822 |
| MDN1 | ENST00000428876 | 2 | 205 | 1 | 502 | 0.205073971 | 1 | 1 | 4.884989822 |
| MACF1 | ENST00000539005 | 2 | 205 | 1 | 502 | 0.205073971 | 1 | 1 | 4.884989822 |
| MACF1 | ENST00000545844 | 2 | 205 | 1 | 502 | 0.205073971 | 1 | 1 | 4.884989822 |
| HIVEP2 | ENST00000012134 | 1 | 206 | 0 | 503 | 0.291549296 | 1 | 1 | Inf |
| TNS1 | ENST00000171887 | 1 | 206 | 0 | 503 | 0.291549296 | 1 | 1 | Inf |

<sup>a</sup>TRID = Transcript ID, <sup>b</sup>Count of subjects with CH or homozygous mutations, <sup>c</sup>1kg = 1000 genomes, <sup>d</sup>bh = Benjamini Hochberg correction, <sup>e</sup>bonf = Bonferroni correction, <sup>f</sup>OR = Odds ratio

**Supplementary Table 5.** Epi4k and 1000 genomes CH and homozygous counts and p-values, EU ancestry, 0.5%MAF, top 50.

| Gene | TRID<sup>a</sup> | Epi4k<br>0.5%<br>yes<sup>b</sup> | Epi4k<br>0.5%<br>no | 1kg<sup>c</sup><br>0.5%<br>yes | 1kg<br>1%<br>no | pval | Pval<br>bh<sup>d</sup> | Pval<br>bonf<sup>e</sup> | OR<sup>f</sup> |
| --- | --- | --- | --- | --- | --- | --- | --- | --- | --- |
| MACF1 | ENST00000289893 | 3 | 204 | 0 | 503 | 0.024527517 | 1 | 1 | Inf |
| MACF1 | ENST00000372915 | 3 | 204 | 0 | 503 | 0.024527517 | 1 | 1 | Inf |
| MACF1 | ENST00000564288 | 3 | 204 | 0 | 503 | 0.024527517 | 1 | 1 | Inf |
| MACF1 | ENST00000567887 | 3 | 204 | 0 | 503 | 0.024527517 | 1 | 1 | Inf |
| OBSCN | ENST00000366707 | 4 | 203 | 2 | 501 | 0.063150347 | 1 | 1 | 4.923199244 |
| MACF1 | ENST00000317713 | 2 | 205 | 0 | 503 | 0.084709668 | 1 | 1 | Inf |
| ABCC11 | ENST00000353782 | 2 | 205 | 0 | 503 | 0.084709668 | 1 | 1 | Inf |
| ABCC11 | ENST00000356608 | 2 | 205 | 0 | 503 | 0.084709668 | 1 | 1 | Inf |
| MACF1 | ENST00000361689 | 2 | 205 | 0 | 503 | 0.084709668 | 1 | 1 | Inf |
| MACF1 | ENST00000372925 | 2 | 205 | 0 | 503 | 0.084709668 | 1 | 1 | Inf |
| PRTG | ENST00000389286 | 2 | 205 | 0 | 503 | 0.084709668 | 1 | 1 | Inf |
| ABCC11 | ENST00000394747 | 2 | 205 | 0 | 503 | 0.084709668 | 1 | 1 | Inf |
| ABCC11 | ENST00000394748 | 2 | 205 | 0 | 503 | 0.084709668 | 1 | 1 | Inf |
| OBSCN | ENST00000441106 | 2 | 205 | 0 | 503 | 0.084709668 | 1 | 1 | Inf |
| MACF1 | ENST00000539005 | 2 | 205 | 0 | 503 | 0.084709668 | 1 | 1 | Inf |
| MACF1 | ENST00000545844 | 2 | 205 | 0 | 503 | 0.084709668 | 1 | 1 | Inf |
| MYO15A | ENST00000205890 | 2 | 205 | 1 | 502 | 0.205073971 | 1 | 1 | 4.884989822 |
| DNAH2 | ENST00000389173 | 2 | 205 | 1 | 502 | 0.205073971 | 1 | 1 | 4.884989822 |
| PKD1L2 | ENST00000525539 | 2 | 205 | 1 | 502 | 0.205073971 | 1 | 1 | 4.884989822 |
| PKD1L2 | ENST00000533478 | 2 | 205 | 1 | 502 | 0.205073971 | 1 | 1 | 4.884989822 |
| DNAH2 | ENST00000572933 | 2 | 205 | 1 | 502 | 0.205073971 | 1 | 1 | 4.884989822 |
| TTN | ENST00000342992 | 17 | 190 | 28 | 475 | 0.234492 | 1 | 1 | 1.516901915 |
| HIVEP2 | ENST00000012134 | 1 | 206 | 0 | 503 | 0.291549296 | 1 | 1 | Inf |
| TNS1 | ENST00000171887 | 1 | 206 | 0 | 503 | 0.291549296 | 1 | 1 | Inf |
| CEACAM21 | ENST00000187608 | 1 | 206 | 0 | 503 | 0.291549296 | 1 | 1 | Inf |
| TGM6 | ENST00000202625 | 1 | 206 | 0 | 503 | 0.291549296 | 1 | 1 | Inf |
| LAMB1 | ENST00000222399 | 1 | 206 | 0 | 503 | 0.291549296 | 1 | 1 | Inf |
| TFR2 | ENST00000223051 | 1 | 206 | 0 | 503 | 0.291549296 | 1 | 1 | Inf |
| DHDDS | ENST00000236342 | 1 | 206 | 0 | 503 | 0.291549296 | 1 | 1 | Inf |
| CCDC92 | ENST00000238156 | 1 | 206 | 0 | 503 | 0.291549296 | 1 | 1 | Inf |
| TICAM1 | ENST00000248244 | 1 | 206 | 0 | 503 | 0.291549296 | 1 | 1 | Inf |
| VIL1 | ENST00000248444 | 1 | 206 | 0 | 503 | 0.291549296 | 1 | 1 | Inf |
| E2F8 | ENST00000250024 | 1 | 206 | 0 | 503 | 0.291549296 | 1 | 1 | Inf |
| DHX29 | ENST00000251636 | 1 | 206 | 0 | 503 | 0.291549296 | 1 | 1 | Inf |
| KRT12 | ENST00000251643 | 1 | 206 | 0 | 503 | 0.291549296 | 1 | 1 | Inf |
| PLXNA1 | ENST00000251772 | 1 | 206 | 0 | 503 | 0.291549296 | 1 | 1 | Inf |
| AKAP12 | ENST00000253332 | 1 | 206 | 0 | 503 | 0.291549296 | 1 | 1 | Inf |
| DNHD1 | ENST00000254579 | 1 | 206 | 0 | 503 | 0.291549296 | 1 | 1 | Inf |
| LPIN1 | ENST00000256720 | 1 | 206 | 0 | 503 | 0.291549296 | 1 | 1 | Inf |
| RABEPK | ENST00000259460 | 1 | 206 | 0 | 503 | 0.291549296 | 1 | 1 | Inf |
| ARHGAP20 | ENST00000260283 | 1 | 206 | 0 | 503 | 0.291549296 | 1 | 1 | Inf |
| TRPV4 | ENST00000261740 | 1 | 206 | 0 | 503 | 0.291549296 | 1 | 1 | Inf |
| NUP153 | ENST00000262077 | 1 | 206 | 0 | 503 | 0.291549296 | 1 | 1 | Inf |
| FBN2 | ENST00000262464 | 1 | 206 | 0 | 503 | 0.291549296 | 1 | 1 | Inf |
| PDE4C | ENST00000262805 | 1 | 206 | 0 | 503 | 0.291549296 | 1 | 1 | Inf |
| KIF18A | ENST00000263181 | 1 | 206 | 0 | 503 | 0.291549296 | 1 | 1 | Inf |
| CLTCL1 | ENST00000263200 | 1 | 206 | 0 | 503 | 0.291549296 | 1 | 1 | Inf |
| LRP2 | ENST00000263816 | 1 | 206 | 0 | 503 | 0.291549296 | 1 | 1 | Inf |
| TECTA | ENST00000264037 | 1 | 206 | 0 | 503 | 0.291549296 | 1 | 1 | Inf |

<sup>a</sup>TRID = Transcript ID, <sup>b</sup>Count of subjects with CH or homozygous mutations, <sup>c</sup>1kg = 1000 genomes, <sup>d</sup>bh = Benjamini Hochberg correction, <sup>e</sup>bonf = Bonferroni correction, <sup>f</sup>OR = Odds ratio

**Supplementary Dataset 1** contains counts of individuals from the Epi4k and 1000 genomes dataset with compound heterozygous or homozygous mutations in each transcript. Results files with counts and p-values for all transcripts are included as well. Dataset S1 is available for viewing on dropbox @ https://www.dropbox.com/sh/yu46zg8bjsafgt9/AACVgJWAj-26mrgsLivJublfa?dl=0

## References

Allache, R., et al. (2012). “Role of the planar cell polarity gene CELSR1 in neural tube defects and caudal agenesis.” Birth Defects Res A Clin Mol Teratol 94(3): 176–181.

Amrutkar, C. and R. M. Riel-Romero (2018). Lennox Gastaut Syndrome. StatPearls. Treasure Island (FL).

Auton A Fau - Brooks, L. D., Brooks Ld Fau - Durbin, R. M., Durbin Rm Fau - Garrison, E. P., Garrison Ep Fau - Kang, H. M., Kang Hm Fau - Korbel, J. O., Korbel Jo Fau - Marchini, J. L.,…Abecasis, G. R. A global reference for human genetic variation. (1476-4687 (Electronic)). doi:D - NLM: NIHMS753481

Bomba, L., Walter, K., & Soranzo, N. (2017). The impact of rare and low-frequency genetic variants in common disease. Genome Biol, 18(1), 77. doi:10.1186/s13059-017-1212-4

Callen, D. F., Ricciardelli, C., Butler, M., Stapleton, A., Stahl, J., Kench, J. G.,…Holm, R. (2010). Co-expression of the androgen receptor and the transcription factor ZNF652 is related to prostate cancer outcome. Oncol Rep, 23(4), 1045–1052.

Cingolani, P., Platts, A., Wang le, L., Coon, M., Nguyen, T., Wang, L.,…Ruden, D. M. (2012). A program for annotating and predicting the effects of single nucleotide polymorphisms, SnpEff: SNPs in the genome of Drosophila melanogaster strain w1118; iso-2; iso-3. Fly (Austin), 6(2), 80–92. doi:10.4161/fly.19695

Clifford, G. M., Rana, R. K., Franceschi, S., Smith, J. S., Gough, G., & Pimenta, J. M. (2005). Human papillomavirus genotype distribution in low-grade cervical lesions: comparison by geographic region and with cervical cancer. Cancer Epidemiol Biomarkers Prev, 14(5), 1157–1164. doi:10.1158/1055-9965.EPI-04-0812

Cornet, I., Gheit, T., Franceschi, S., Vignat, J., Burk, R. D., Sylla, B. S.,…Group, I. H. V. S. (2012). Human papillomavirus type 16 genetic variants: phylogeny and classification based on E6 and LCR. J Virol, 86(12), 6855–6861. doi:10.1128/JVI.00483-12

Correa, R. G., Krajewska, M., Ware, C. F., Gerlic, M., & Reed, J. C. (2014). The NLR-related protein NWD1 is associated with prostate cancer and modulates androgen receptor signaling. Oncotarget, 5(6), 1666–1682. doi:10.18632/oncotarget.1850

Covanis, A. (2012). “Epileptic encephalopathies (including severe epilepsy syndromes).” Epilepsia 53 Suppl 4: 114–126.

Cox, A. J., Darbro, B. W., Laxer, R. M., Velez, G., Bing, X., Finer, A. L.,…Ferguson, P. J. (2017). Recessive coding and regulatory mutations in FBLIM1 underlie the pathogenesis of chronic recurrent multifocal osteomyelitis (CRMO). PLoS One, 12(3), e0169687. doi:10.1371/journal.pone.0169687

DeScipio, C., et al. (2012). “Subtelomeric deletion of chromosome 10p15.3: clinical findings and molecular cytogenetic characterization.” Am J Med Genet A 158A(9): 2152–2161.

Dobyns, W. B., et al. (2018). “MACF1 Mutations Encoding Highly Conserved Zinc-Binding Residues of the GAR Domain Cause Defects in Neuronal Migration and Axon Guidance.” Am J Hum Genet.

Dolinsky, T. J., Nielsen, J. E., McCammon, J. A., & Baker, N. A. (2004). PDB2PQR: an automated pipeline for the setup of Poisson-Boltzmann electrostatics calculations. Nucleic Acids Res, 32(Web Server issue), W665–667. doi:10.1093/nar/gkh381

Edwards, A., Hammond, H. A., Jin, L., Caskey, C. T., & Chakraborty, R. (1992). Genetic variation at five trimeric and tetrameric tandem repeat loci in four human population groups. Genomics, 12(2), 241–253.

Epi, K. C., Epilepsy Phenome/Genome, P., Allen, A. S., Berkovic, S. F., Cossette, P., Delanty, N.,…Winawer, M. R. (2013). De novo mutations in epileptic encephalopathies. Nature, 501(7466), 217–221. doi:10.1038/nature12439

Genomes Project, C., Abecasis, G. R., Altshuler, D., Auton, A., Brooks, L. D., Durbin, R. M.,…McVean, G. A. (2010). A map of human genome variation from population-scale sequencing. Nature, 467(7319), 1061–1073. doi:10.1038/nature09534

Genomes Project, C., Abecasis, G. R., Auton, A., Brooks, L. D., DePristo, M. A., Durbin, R. M.,…McVean, G. A. (2012). An integrated map of genetic variation from 1,092 human genomes. Nature, 491(7422), 56–65. doi:10.1038/nature11632

Genomes Project, C., Auton, A., Brooks, L. D., Durbin, R. M., Garrison, E. P., Kang, H. M.,…Abecasis, G. R. (2015). A global reference for human genetic variation. Nature, 526(7571), 68–74. doi:10.1038/nature15393

Giovannucci, E., Stampfer, M. J., Krithivas, K., Brown, M., Dahl, D., Brufsky, A.,…Kantoff, P. W. (1997). The CAG repeat within the androgen receptor gene and its relationship to prostate cancer. Proc Natl Acad Sci U S A, 94(7), 3320–3323.

Goncalves, C. M., Castro, M. A., Henriques, T., Oliveira, M. I., Pinheiro, H. C., Oliveira, C.,…Carmo, A. M. (2009). Molecular cloning and analysis of SSc5D, a new member of the scavenger receptor cysteine-rich superfamily. Mol Immunol, 46(13), 2585–2596. doi:10.1016/j.molimm.2009.05.006

Haiman, C. A., Chen, G. K., Blot, W. J., Strom, S. S., Berndt, S. I., Kittles, R. A.,…Henderson, B. E. (2011). Genome-wide association study of prostate cancer in men of African ancestry identifies a susceptibility locus at 17q21. Nat Genet, 43(6), 570–573. doi:10.1038/ng.839

Ho, L., Chan, S. Y., Burk, R. D., Das, B. C., Fujinaga, K., Icenogle, J. P.,…et al. (1993). The genetic drift of human papillomavirus type 16 is a means of reconstructing prehistoric viral spread and the movement of ancient human populations. J Virol, 67(11), 6413–6423.

Jakovcevski, I., Miljkovic, D., Schachner, M., & Andjus, P. R. (2013). Tenascins and inflammation in disorders of the nervous system. Amino Acids, 44(4), 1115–1127. doi:10.1007/s00726-012-1446-0

Jelen, M. M., Chen, Z., Kocjan, B. J., Burt, F. J., Chan, P. K., Chouhy, D.,…Poljak, M. (2014). Global genomic diversity of human papillomavirus 6 based on 724 isolates and 190 complete genome sequences. J Virol, 88(13), 7307–7316. doi:10.1128/JVI.00621-14

Jelen, M. M., Chen, Z., Kocjan, B. J., Hosnjak, L., Burt, F. J., Chan, P. K.,…Poljak, M. (2016). Global Genomic Diversity of Human Papillomavirus 11 Based on 433 Isolates and 78 Complete Genome Sequences. J Virol, 90(11), 5503–5513. doi:10.1128/JVI.03149-15

Konecny, R., Baker, N. A., & McCammon, J. A. (2012). iAPBS: a programming interface to Adaptive Poisson-Boltzmann Solver (APBS). Comput Sci Discov, 5(1). doi:10.1088/1749-4699/5/1/015005

Kosmicki, J. A., Churchhouse, C. L., Rivas, M. A., & Neale, B. M. (2016). Discovery of rare variants for complex phenotypes. Hum Genet, 135(6), 625–634. doi:10.1007/s00439-016-1679-1

Kossoff, E. H. (2010). “Infantile spasms.” Neurologist 16(2): 69–75.

Lauring, A. S., Frydman, J., & Andino, R. (2013). The role of mutational robustness in RNA virus evolution. Nat Rev Microbiol, 11(5), 327–336. doi:10.1038/nrmicro3003

Lek, M., Karczewski, K., Minikel, E., Samocha, K., Banks, E., Fennell, T.,…MacArthur, D. (2015). Analysis of protein-coding genetic variation in 60,706 humans. bioRxiv. doi:10.1101/030338

Lek, M., Karczewski, K. J., Minikel, E. V., Samocha, K. E., Banks, E., Fennell, T.,…Exome Aggregation, C. (2016). Analysis of protein-coding genetic variation in 60,706 humans. Nature, 536(7616), 285–291. doi:10.1038/nature19057

Li, Y., Vinckenbosch, N., Tian, G., Huerta-Sanchez, E., Jiang, T., Jiang, H.,…Wang, J. (2010). Resequencing of 200 human exomes identifies an excess of low-frequency non-synonymous coding variants. Nat Genet, 42(11), 969–972. doi:10.1038/ng.680

Lin, M. S., et al. (2017). “alpha2-glycine receptors modulate adult hippocampal neurogenesis and spatial memory.” Dev Neurobiol 77(12): 1430–1441.

Martinez, V. G., Moestrup, S. K., Holmskov, U., Mollenhauer, J., & Lozano, F. (2011). The conserved scavenger receptor cysteine-rich superfamily in therapy and diagnosis. Pharmacol Rev, 63(4), 967–1000. doi:10.1124/pr.111.004523

McKenna, A., Hanna, M., Banks, E., Sivachenko, A., Cibulskis, K., Kernytsky, A.,…DePristo, M. A. (2010). The Genome Analysis Toolkit: a MapReduce framework for analyzing next-generation DNA sequencing data. Genome Res, 20(9), 1297–1303. doi:10.1101/gr.107524.110

Mercado-Gomez, O., Landgrave-Gomez, J., Arriaga-Avila, V., Nebreda-Corona, A., & Guevara-Guzman, R. (2014). Role of TGF-beta signaling pathway on Tenascin C protein upregulation in a pilocarpine seizure model. Epilepsy Res, 108(10), 1694–1704. doi:10.1016/j.eplepsyres.2014.09.019

Miro-Julia, C., Rosello, S., Martinez, V. G., Fink, D. R., Escoda-Ferran, C., Padilla, O.,…Lozano, F. (2011). Molecular and functional characterization of mouse S5D-SRCRB: a new group B member of the scavenger receptor cysteine-rich superfamily. J Immunol, 186(4), 2344–2354. doi:10.4049/jimmunol.1000840

Miwa, H., Go, M. F., & Sato, N. (2002). H. pylori and gastric cancer: the Asian enigma. Am J Gastroenterol, 97(5), 1106–1112. doi:10.1111/j.1572-0241.2002.05663.x

Moffat, J. J., Ka, M., Jung, E. M., Smith, A. L., & Kim, W. Y. (2017). The role of MACF1 in nervous system development and maintenance. Semin Cell Dev Biol, 69, 9–17. doi:10.1016/j.semcdb.2017.05.020

Moshfegh, Y., Velez, G., Li, Y., Bassuk, A. G., Mahajan, V. B., & Tsang, S. H. (2016). BESTROPHIN1 mutations cause defective chloride conductance in patient stem cell-derived RPE. Hum Mol Genet, 25(13), 2672–2680. doi:10.1093/hmg/ddw126

O’Brien, T. G., Guo, Y., Visvanathan, K., Sciulli, J., McLaine, M., Helzlsouer, K. J., & Watkins-Bruner, D. (2004). Differences in ornithine decarboxylase and androgen receptor allele frequencies among ethnic groups. Mol Carcinog, 41(2), 120–123. doi:10.1002/mc.20047

Oliveros, J. C. (2015). Venny. An interactive tool for comparing lists with Venn’s diagrams.

Peprah, E., Xu, H., Tekola-Ayele, F., & Royal, C. D. (2015). Genome-wide association studies in Africans and African Americans: expanding the framework of the genomics of human traits and disease. Public Health Genomics, 18(1), 40–51. doi:10.1159/000367962

Pilorge, M., et al. (2016). “Genetic and functional analyses demonstrate a role for abnormal glycinergic signaling in autism.” Mol Psychiatry 21(7): 936–945.

Polley, S., Louzada, S., Forni, D., Sironi, M., Balaskas, T., Hains, D. S.,…Hollox, E. J. (2015). Evolution of the rapidly mutating human salivary agglutinin gene (DMBT1) and population subsistence strategy. Proc Natl Acad Sci U S A, 112(16), 5105–5110. doi:10.1073/pnas.1416531112

Rand, K. A., Rohland, N., Tandon, A., Stram, A., Sheng, X., Do, R.,…Haiman, C. A. (2016). Whole-exome sequencing of over 4100 men of African ancestry and prostate cancer risk. Hum Mol Genet, 25(2), 371–381. doi:10.1093/hmg/ddv462

Robinson, A., et al. (2012). “Mutations in the planar cell polarity genes CELSR1 and SCRIB are associated with the severe neural tube defect craniorachischisis.” Hum Mutat 33(2): 440–447.

Salter, J. D., Bennett, R. P., & Smith, H. C. (2016). The APOBEC Protein Family: United by Structure, Divergent in Function. Trends Biochem Sci, 41(7), 578–594. doi:10.1016/j.tibs.2016.05.001

Sanjak, J. S., Long, A. D., & Thornton, K. R. (2017). A Model of Compound Heterozygous, Loss-of-Function Alleles Is Broadly Consistent with Observations from Complex-Disease GWAS Datasets. PLoS Genet, 13(1), e1006573. doi:10.1371/journal.pgen.1006573

Stelzer, G., Inger, A., Olender, T., Iny-Stein, T., Dalah, I., Harel, A.,…Lancet, D. (2009). GeneDecks: paralog hunting and gene-set distillation with GeneCards annotation. OMICS, 13(6), 477–487. doi:10.1089/omi.2009.0069

Toral, M. A., Velez, G., Boudreault, K., Schaefer, K. A., Xu, Y., Saffra, N.,…Mahajan, V. B. (2017). Structural modeling of a novel SLC38A8 mutation that causes foveal hypoplasia. Mol Genet Genomic Med, 5(3), 202–209. doi:10.1002/mgg3.266

Toyoda, R., Nakamura, H., & Watanabe, Y. (2005). Identification of protogenin, a novel immunoglobulin superfamily gene expressed during early chick embryogenesis. Gene Expr Patterns, 5(6), 778–785. doi:10.1016/j.modgep.2005.04.001

Vesque, C., Anselme, I., Couve, E., Charnay, P., & Schneider-Maunoury, S. (2006). Cloning of vertebrate Protogenin (Prtg) and comparative expression analysis during axis elongation. Dev Dyn, 235(10), 2836–2844. doi:10.1002/dvdy.20898

Vieira, V. C., Leonard, B., White, E. A., Starrett, G. J., Temiz, N. A., Lorenz, L. D.,…Harris, R. S. (2014). Human papillomavirus E6 triggers upregulation of the antiviral and cancer genomic DNA deaminase APOBEC3B. MBio, 5(6). doi:10.1128/mBio.02234-14

W, D. The PyMOL Molecular Graphics System.

Wang, C., Zhan, X., Bragg-Gresham, J., Kang, H. M., Stambolian, D., Chew, E. Y.,…Abecasis, G. R. Ancestry estimation and control of population stratification for sequence-based association studies. (1546-1718 (Electronic)). doi:D - NLM: NIHMS588246

D - NLM: PMC4084909 EDAT-2014/03/19 06:00 MHDA-2014/05/20 06:00 CRDT-2014/03/18 06:00 PHST-2013/08/08 [received] PHST-2014/02/21 [accepted] AID - ng.2924 [pii] AID - 10.1038/ng.2924 [doi] PST - ppublish

Warren, C. J., Westrich, J. A., Doorslaer, K. V., & Pyeon, D. (2017). Roles of APOBEC3A and APOBEC3B in Human Papillomavirus Infection and Disease Progression. Viruses, 9(8). doi:10.3390/v9080233

Webb, B., & Sali, A. (2016). Comparative Protein Structure Modeling Using MODELLER. Curr Protoc Protein Sci, 86, 2 9 1–2 9 37. doi:10.1002/cpps.20

Zaidi, S. F. (2016). Helicobacter pylori associated Asian enigma: Does diet deserve distinction? World J Gastrointest Oncol, 8(4), 341–350. doi:10.4251/wjgo.v8.i4.341

Zarrei, M., et al. (2018). “De novo and rare inherited copy-number variations in the hemiplegic form of cerebral palsy.” Genet Med 20(2): 172–180.

Zhang, Y., et al. (2017). “Structure-Function Analysis of the GlyR alpha2 Subunit Autism Mutation p.R323L Reveals a Gain-of-Function.” Front Mol Neurosci 10: 158.

Zhong, K., Karssen, L. C., Kayser, M., & Liu, F. (2016). CollapsABEL: an R library for detecting compound heterozygote alleles in genome-wide association studies. BMC Bioinformatics, 17, 156. doi:10.1186/s12859-016-1006-9

